# Engineered lactobacilli display anti-biofilm and growth suppressing activities against *Pseudomonas aeruginosa*

**DOI:** 10.1101/2020.04.08.032946

**Authors:** Todd C. Chappell, Nikhil U. Nair

**Affiliations:** Department of Chemical & Biological Engineering, Tufts University

**Keywords:** Biofilm, Bacteriotherapy, Lactobacillus, Antimicrobial, Probiotic, *P. aeruginosa*

## Abstract

Biofilms are an emerging target for new therapeutics in the effort to address the continued increase in resistance and tolerance to traditional antimicrobials. In particular, the distinct nature of the biofilm growth state often means that traditional antimicrobials, developed to combat planktonic cells, are ineffective. Biofilm treatments are designed to both reduce pathogen load at an infection site and decrease the development of resistance by rendering the embedded organisms more susceptible to treatment at lower antimicrobial concentrations. In this work, we developed a new antimicrobial treatment modality by characterizing the natural capacity of two lactobacilli, *L. plantarum* and *L. rhamnosus*, to inhibit *P. aeruginosa* growth, biofilm formation, and biofilm viability. We further engineered these lactic acid bacteria (LAB) to secrete enzymes known to degrade *P. aeruginosa* biofilms and show that our best performing engineered LAB, secreting a pathogen-derived enzyme (PelA_hyd_), degrades up to 85 % of *P. aeruginosa* biofilm.

## 1. Introduction

As an important virulence factor for pathogenic microbes, biofilms are associated with an expanding array of pathologies, including various airway, gastrointestinal, and ocular infections, endocarditis, periodontitis, osteomyelitis, cystitis, and chronic wounds^1–7^. Biofilms represent a distinct growth state, morphologically distinguished by bacteria residing within a self-produced matrix of extracellular polymeric substances (EPS), that may include proteins, extracellular DNA (eDNA), polysaccharides, and lipids^8^. Within the biofilm, isogenic cells exhibit phenotypic diversity that is driven by the discrete microenvironments created by metabolite, ion, gas, and antimicrobial diffusion gradients into and out of the biofilm. Biomedically, this phenotypic diversity manifests as distinct tolerances or resistances to traditional antimicrobials, as well as the host immune system^9,10^. Additionally, biofilms stabilize surface colonization and are frequently less susceptible to traditional methods for surface decontamination, exacerbating the recalcitrance to treatment. Thus, clearance of mature biofilms is an essential component for the successful resolution of numerous infections, especially those that are chronic or recurrent in nature.

Invasive burn wounds and chronic wounds, or wounds that fail to progress though the later stages of the normal healing process, are commonly contaminated or colonized by a multitude of biofilm-forming organisms. Standard treatments for these wound types include nanocrystalline silver, silver sulphadiazine, iodine, or topical antibiotics. However, these treatments are often ineffective at reducing wound infection, add unnecessary expense, and/or inhibit the healing process^11–14^. Further, extensive use of these treatments has bred a large population of multi-drug resistant microbes for which new treatments that target both planktonic and biofilms cells are necessary.

A popular biofilm targeting strategy is the enzymatic degradation of biofilm polymer(s) to decrease surface adhesion and return the entrained bacteria to a more treatable phenotype. Rapid advancement in synthetic biology and probiotic therapies have led to interest in developing engineered bacteriotherapies or live biotherapeutic products. These “smart”, bacteria-based therapeutic delivery vectors provide sustained delivery of the therapeutic and dynamically respond to environmental signals, while retaining their innate probiotic qualities^15–18^. Recent examples of bacteriotherapies include the delivery of enzymes, antimicrobials, metabolites, or anti-inflammatory proteins to combat metabolic deficiencies, tumors, inflammation, biofilms and infections^15,19–25^. In this study, we construct and assess the utility of genetically engineered probiotic bacteria as anti-biofilm and antimicrobial agents against the common wound pathogen *Pseudomonas aeruginosa*.

We selected lactic acid bacteria (LAB) as the chassis strains for the bacteriotherapy due to their broad-spectrum antimicrobial and wound healing capacities, genetic tractability, and well characterized expression systems for the production and secretion of heterologous proteins. Furthermore, Several LAB have been shown to impair the growth of drug resistant *P. aeruginosa* clinical isolates^26^. More specifically, *Lactobacillus plantarum* and *Lactobacillus rhamnosus*, the species used in this work, enhance the outcome of mouse *P. aeruginosa* infection models and increase epithelial migration, and are equally as effective as current treatments when applied to human burn wounds^27–30^. We add to this body of evidence, showing that *L. plantarum* WCFS1 and *L. rhamnosus* (LGG) are effective inhibitors of PA14 planktonic growth, while also inhibiting biofilm formation and the viability of PA14 biofilm embedded cells (biofilm viability). We further increase the usefulness of *L. plantarum* and LGG by engineering them to secrete enzymes known to degrade PA14 biofilms and demonstrate the efficacy of this design for degradation of mature PA14 biofilms.

## 2. Materials and Methods

### 2.1. Bacterial growth and transformation

All strains used in this study are listed in Table S1. *E. coli* strains were grown in LB broth and plated on LB agar, unless stated otherwise. Erythromycin and ampicillin were added to *E. coli* cultures at 200 or 100 µg/mL, respectively. Lactobacilli were grown in De Man, Rogosa and Sharpe (MRS; RPI Corp) broth and plated on MRS agar (1.5 % w/v) plates unless stated otherwise. Erythromycin was added lactobacillus cultures at 5 µg/mL when necessary. All cultures were grown at 37 °C; *E. coli and L. plantarum* cultures were grown shaking (250 rpm) and LGG was grown statically, unless stated otherwise. *E. coli* transformation was performed using MES or TSS competent cells. *L. plantarum* WCFS1 transformation was performed using a method derived from Aukrust and Blom^31^. LGG transformation was performed using the method described in De Keersmaecker et al.^32^.

### 2.2. LAB antimicrobial plate assay

Overnight cultures of LAB were diluted 1000× into 10 mL fresh media and 1 mL aliquots were removed from the culture at the designated times. Following aliquot removal from LAB culture, cells were pelleted at 4000 ×g for 15 min and the resulting supernatant was filtered through 0.22 µm PES filter and frozen at ─20 °C until the following day. The following day, overnight cultures of *P. aeruginosa* were diluted 100× in LB broth and 100 µL was plated on the surface of LB agar. Agar wells were excised from the agar plate and 200 µL of fresh lactobacilli culture or filtered culture supernatant was added to each well. Plates were incubated at 37 °C overnight and inhibition was evaluated qualitatively by inhibition of pathogen growth.

### 2.3. Plasmid construction

All vectors used and constructed in this study are listed in Table S2. *E. coli* TG1 was modified by knockout of *endA* to improve transformation efficiency and plasmid quality of pSIP411-derived vectors. DH5α was used to propagate all pSIP401-derived plasmids. All DNA oligos were ordered from Eurofins or Genewiz and sequences are given in Table S3. Gene knockout and verification in *E. coli* TG1 was performed using λRed recombineering^33^ as described previously using primers 17-20^34^. Variants of the pSIP401 and pSIP411 plasmids with inserts containing the Lp_3050 secretion signal, 6× histidine tag, thrombin cleavage site, and multiple cloning site (MCS) were constructed using primers 1, 2, 3, 4, 5 (Table S3). The inserts for these constructs were generated by overlap extension PCR. The product and vectors (pSIP401 and pSIP411) were digested with *Bgl*II and *Pml*I to construct pTCC200 and pTCC210. The *nucA* gene was amplified from genomic DNA prepared from *Staphylococcus aureus* UAMS1 using primers 6 and 7. The *engZ* gene was amplified from genomic DNA of *Clostridium cellulovorans* (purchased from DSMZ) using primers 8 and 9. Amplified DNA fragments containing *nucA* or *engZ* were digested with *Sal*I and *Pml*I for insertion into the same digested pTCC200. The gene for PelA_hyd_ was amplified from genomic DNA prepared from PA14 using primers 10 and 11. Plasmid pTCC210 was amplified using primers 12 and 13 and combined with the PelA_hyd_ fragment using NEBuilder® HiFi DNA Assembly. All enzymes were purchased from New England Biolabs (NEB, Ipswich, MA). All cloning inserts were amplified using Phusion® DNA polymerase. All inserts in modified plasmids were verified by colony PCR and sequenced by Eurofins Genomics LLC (Louisville, KY) or Genewiz, Inc. (Cambridge, MA) using primers 14,15, and 16.

### 2.4. Liquid culture biofilm formation

*P. aeruginosa* PA14 biofilms were grown by diluting a 24 h culture 200× into salt-free LB (sfLB). For biofilm inhibition studies, PA14 was also diluted 200× into the supernatants from 24 h cultures of *L. plantarum* and LGG, and this culture was subsequently serially diluted into new sfLB PA14 subculture to maintain consistent cell density. The new PA14 cultures were dispensed in 150 μL aliquots into wells of white Lumitrac high-bind 96-well microplates (Greiner Bio-One, Monroe, NC). Wells at the plate edge were filled with water and only interior wells were used for biofilm formation. Microplate lids were sealed with parafilm and incubated for 24 h at 30 °C without shaking. Biofilms biomass was then quantified or incubated further to measure treatment efficacy.

### 2.5. Biofilm quantification

The biofilm biomass was measured by staining adherent cells with crystal violet (CV). Wash steps were performed using low pipette flow rates to prevent removal of adherent cells. Biofilms grown in liquid cultures as described above were washed 2× with 250 µL PBS to remove non-adherent cells. 250 µL of aqueous 0.1 % CV was added to each well, and plates were incubated for 15 min. Following incubation, plates were inverted and washed 4× with 300 µL phosphate-buffered saline (PBS; 8 g/L NaCl, 0.2 g/L KCl, 10 mM Na_2_HPO_4_, 1.8 mM KH_2_PO_4_). Plates were dried at 37 °C. De-staining was performed by addition of 300 µL of 4:1 ethanol:acetone solution. After 15 – 20 min, 200 µL of the solubilized CV solution was transferred to a new 96-well microplate, and the absorbance at 570 nm wavelength was measured (Spectramax^®^ M3, Molecular Devices, San Jose, CA).

### 2.6. Enzyme induction and secretion validation

Protein expression in *L. plantarum* and LGG was performed by growing overnight cultures in MRS and sub-culturing to OD_600_ of 0.05 in BHI supplemented with 0.5 % (w/v) glucose. Cultures were grown until OD_600_ 0.2 – 0.3, pelleted, and induced by resuspending in fresh 2× BHI 0.5% glucose with 200 ng/mL IP-673. Induced cultures were grown for approximately 5 h at 30 °C. For induced supernatants, cells were pelleted by centrifugation 4500 ×g for 10 min, and the supernatant was filtered through a 0.22 µm PES filter. For SDS-PAGE, 10 µL of 4× loading buffer was added to 30 µL LAB supernatant and heated at 95 °C for 5 min. 20 µL of the processed supernatant was loaded per well on a 4 – 12 % Bis-Tris gradient gel. The gel was run in 1× MOPS running buffer at 120 V for approximately 1.5 h and developed using silver stain (Pierce Biotechnology Inc., Rockford, IL). Cellulase activity was evaluated by aliquoting 5 µL of induced cell supernatants on 1.0 % agar plates containing 0.5 % CMC (carboxymethyl cellulose) and incubated overnight at 37 °C. Plates were subsequently incubated with 0.5 % Congo Red (CR) for 10 min. Residual CR was removed by de-staining with 1 M NaCl. DNase activity was assessed by aliquoting 5 µL of induced cell supernatants were spotted on 1.0 % agar plates containing 0.2 % DNA and incubated overnight at 37 °C. Plates were then treated with 1 N HCl to precipitate residual DNA.

### 2.7. Enzymatic degradation of PA14 biofilms by engineered LAB

Efficacy of enzymatic treatment was determined by growing biofilms as described above. The supernatant and nonadherent solids of biofilm cultures were aspirated using a multichannel pipette. 250 µL of induced LAB culture was aliquoted per well, and biofilm microplates were placed on a rocker at room temperature for 1.5 h. LAB cultures were aspirated from wells were the plates were washed and twice with 250 µL of sterile DPBS (2.67 mM KCl, 136.9 mM NaCl, 1.47 mM KH_2_PO_4_, 8.10 mM Na_2_HPO_4_). Remaining biofilm was fixed to plates by drying plates overnight in a 37 °C incubator and quantified using the CV method described previously.

### 2.8. Biofilm viability assay

PA14 biofilms were grown as described above. Filtered LAB supernatants from 24 h LAB cultures were generated as described in the agar-well diffusion assay, and then serially diluted 2× into PBS pH 7.4. The supernatant from the biofilm cultures was removed and 250 µL of diluted LAB supernatants or buffered MRS control was added to each well and the plates were incubated at 37 °C for 24 h. The supernatant was then removed, and the plates were washed 2x with PBS to remove nonadherent cells. 200 µL of LB and 100 µL of XTT solution were added to each well and plates were incubated at 37C for 3 h. Microplates were centrifuged at 3000xg for 10 min and 200 µL of solution was aliquoted into a fresh microplate. The absorbance at 475 nm was taken to determine viability.

## 3. Results

### 3.1. *L. plantarum* and LGG inhibit PA14 growth in a pH-dependent manner

The feasibility of *L. plantarum* and LGG as therapeutic vectors was first analyzed by characterizing their innate capacity to inhibit PA14 growth using an agar-well diffusion assay and dilutions of LAB cultures in a modified MIC assay. The agar-well diffusion assay was used to determine the aeration and duration of LAB culture that maximally inhibited PA14 growth. When *L. plantarum* cultures were shaken in a test tube or flask, growth inhibition of PA14 moderately increased (Fig. S1). However, culture aeration had no impact on LGG growth inhibition of PA14. Early phase *L. plantarum* and LGG cultures (grown ≤ 4 h) and supernatants failed to inhibit PA14 growth, while late-stage (≥ 8 h) cultures and supernatants of both organisms inhibited PA14 growth (Fig. 1A, Fig. S1). 24 h cultures of *L. plantarum* were marginally more inhibitory than those of LGG; yet the supernatants of both LAB exhibited similar growth inhibition against PA14. Generally, we found that PA14 growth inhibition increased with LAB culture duration and cultures were more inhibitory than cell-free supernatants. The pH of 24 h supernatants was 3.8 ─ 3.9 and when we adjusted their pH back up to the starting pH of 6.3, we observed no growth inhibition. To determine if the inhibitory activity was due to pH alone or a factor that was active at low pH, we adjusted the pH of fresh MRS down to that of spent media and evaluated its inhibitory activity against PA14. Decreasing the medium pH increased growth inhibition and when adjusted to pH 3.5, the medium had similar inhibitory activity as that of a 24 h LAB culture of pH 3.8 ─ 3.9.

**Figure 1:**
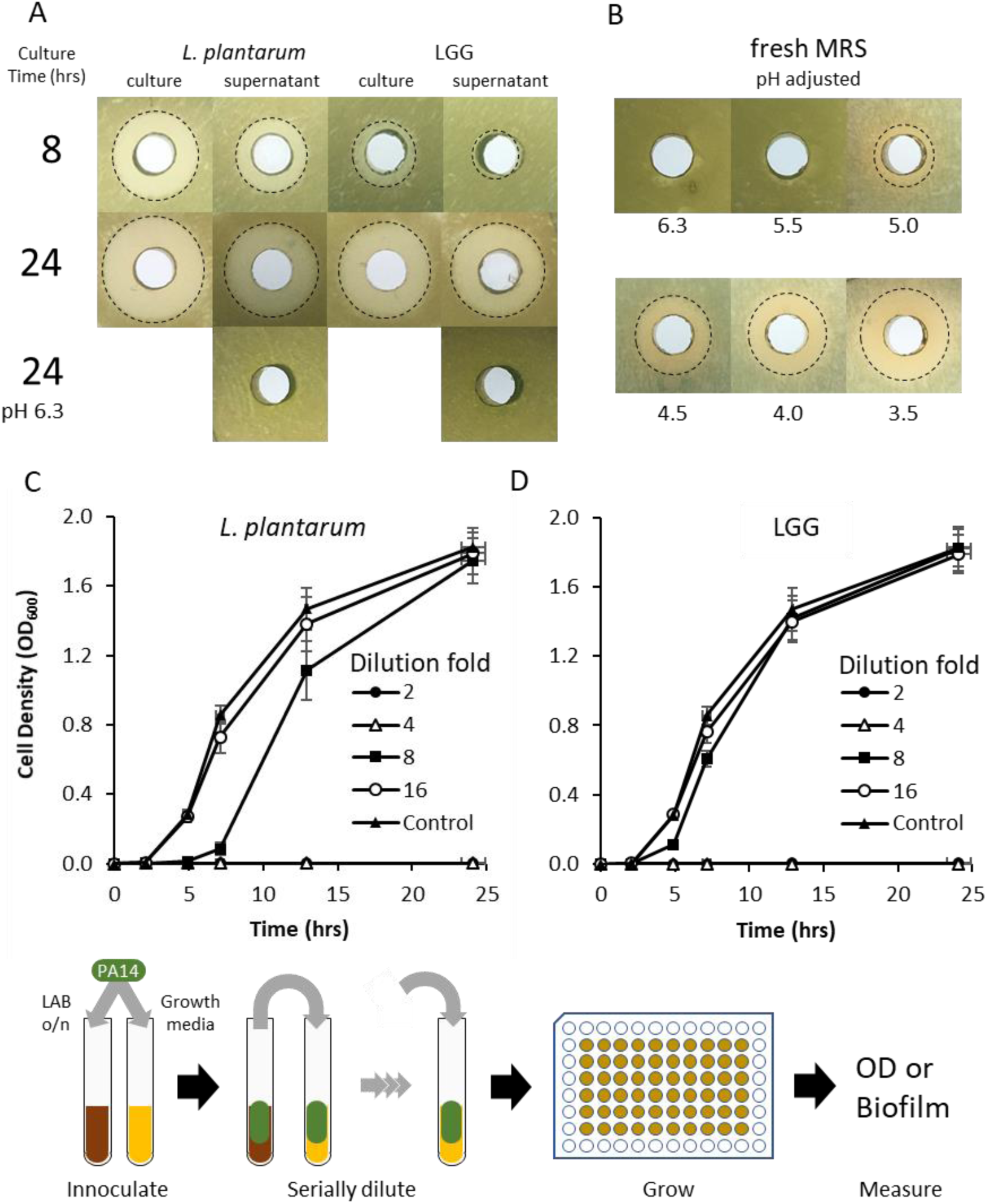
Inhibition of PA14 growth by LAB. (A) Agar well diffusion assay of *L. plantarum* and LGG cultures and supernatants grown in MRS medium. Culture time was either 8 or 24 h. pH adjustment abrogates inhibitor activity of supernatants. (B) Agar well diffusion assay of pH-adjusted fresh MRS medium. The pH of the base-adjusted supernatant and medium is located below the plate image to which it refers. Cultures of planktonic PA14 with (C) *L. plantarum* and (D) LGG supernatants show inhibition at low dilutions factors. Workflow for (C) and (D) is shown: The LAB supernatants were inoculated with PA14, which were serially diluted into fresh PA14 cultures. Each line represents a different dilution factor of the LAB supernatant.

We also used a modified MIC (minimum inhibitory concentration) assay to more quantitatively evaluate the inhibition of PA14 growth by *L. plantarum* and LGG supernatants over time and determine the relative quantity of spent supernatant necessary for bioactivity. Supernatants from 24 h cultures of *L. plantarum* and LGG were diluted to 25 % (i.e., 4× dilution) of the culture volume completely inhibited PA14 growth (Fig. 1C & D). *L. plantarum* supernatant diluted 8×still retained some inhibitory activity, while 8× dilution of LGG supernatants had no inhibitory activity relative to growth medium alone. A dilution of 16×, or greater, of either LAB culture supernatant failed to inhibit PA14 growth.

### 3.2. *L. plantarum* and LGG inhibit PA14 biofilm formation and viability

Having characterized *L. plantarum* and LGG inhibition of planktonic PA14 cells, we also analyzed the impact *L. plantarum* and LGG supernatants had on PA14 biofilm formation and biofilm viability (i.e., viability of cells in the biofilm matrix). We used the modified MIC assay workflow to evaluate the inhibition of PA14 biofilm formation. The supernatants from *L. plantarum* and LGG cultures inhibited PA14 biofilm formation in a concentration dependent manner (Fig. 2A). Only at dilutions greater than 16× did biofilm form at detectable levels. The MRS media control also inhibited PA14 biofilm formation, but only when undiluted and diluted 2-fold.

**Figure 2:**
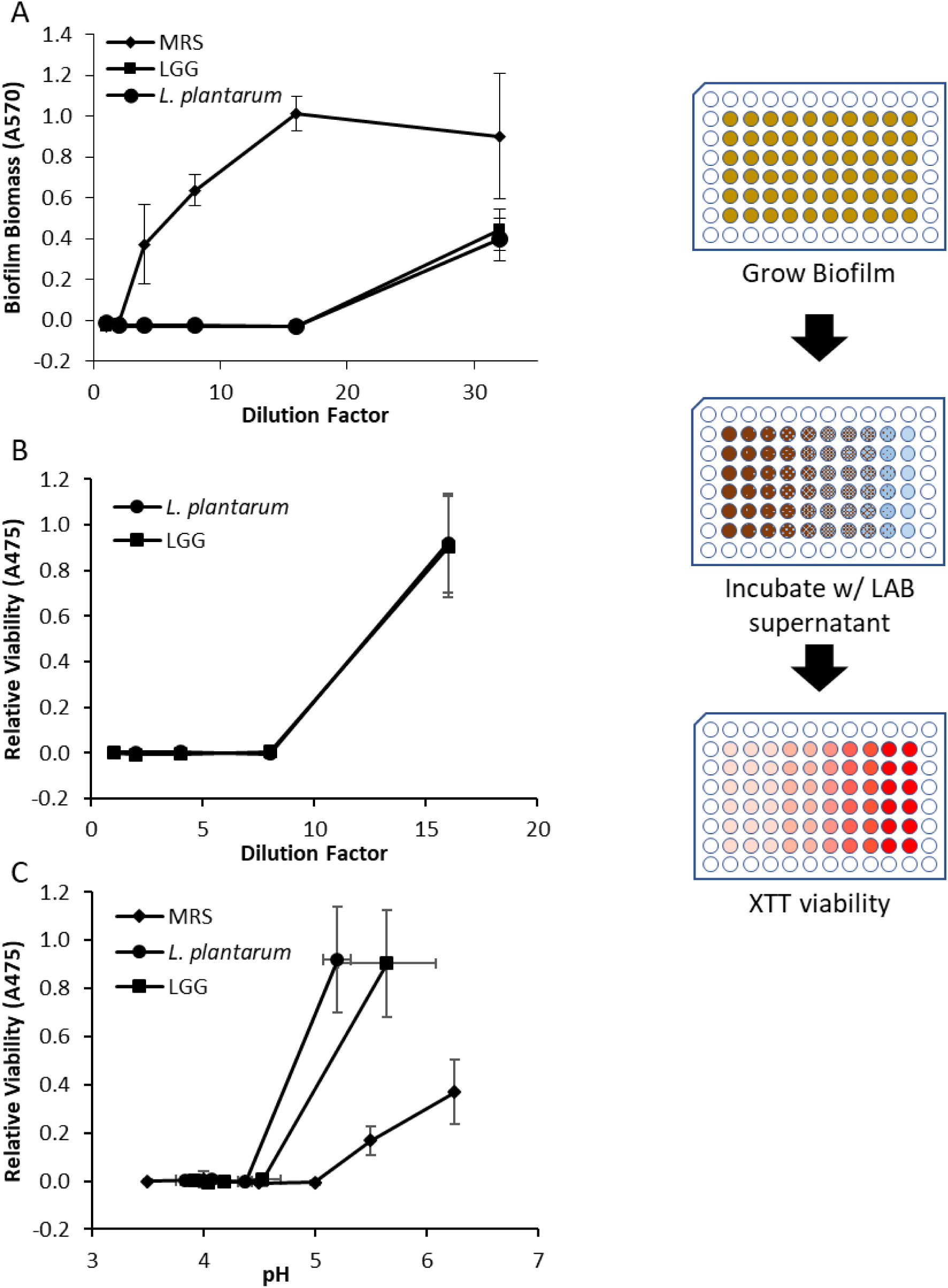
Anti-PA14 biofilm activity of LAB. (A) Inhibition of PA14 biofilm formation by *L. plantarum* and LGG. Workflow for A was similar to that shown in Figure 1. (B) Inhibition of viability of mature PA14 biofilms by *L. plantarum* and LGG. LAB cultures were diluted in PBS. (C) Inhibition of PA14 biofilm viability by *L. plantarum* and LGG compared to pH of culture diluted in PBS or MRS adjusted to a specific pH. Workflow for (B) and (C) is shown.

LAB cell-free supernatants also inhibited the viability of PA14 cells embedded within biofilms, as assessed by XTT dye assay^35^ (Fig. 2B). *L. plantarum* and LGG supernatants diluted by 8×or less were able to inhibit PA14 biofilm viability such that no viable cells could be detected relative to the control. When we plotted the viability against the pH of LAB culture dilutions, we found that the transition to viable biofilms correlates with the increase in pH caused by dilution into PBS (Fig. 2C). Fresh MRS buffered to pH 5 and below completely inhibited biofilm viability—a finding in agreement with our previous finding that the medium has an innate capacity to inhibit PA14 growth when adjusted to a lower pH. Interestingly, MRS adjusted to pH 6.25 and 5.5 also inhibited PA14 biofilm viability more than the diluted LAB cultures of a similar pH, indicating innate anti-PA14 biofilm activity in the MRS. However, we found the major driver of decreased biofilm viability to be low pH. Generally, diluted solutions with a pH ≤ 4.5 were nonviable, while solutions with a pH ≥ 5.2 were viable. We also found that for a given pH, the inhibitory activity of undiluted spent LAB supernatant is more compared to that diluted in PBS.

### 3.3. *L. plantarum* secreted matrix-degrading enzymes disrupt mature PA14 biofilms

*P. aeruginosa* biofilms are predominantly composed of an array of polysaccharides (Alg, Psl, Pel), and eDNA, but the specific composition is dependent upon the genetic background. The strain *P. aeruginosa* PA14, a burn wound isolate, produces biofilms predominantly composed of Pel polysaccharide and eDNA. *P. aeruginosa* biofilms containing these components were previously shown to be sensitive to enzymatic degradation by solutions containing DNase, cellulase ^36,37^, or PelA_hyd_ (the hydrolase domain of PelA) a native enzyme from *P. aeruginosa* that hydrolyzes the Pel polysaccharide to release biofilm cells and transition to planktonic growth^38^. We constructed broad host range LAB expression vectors for secretion of the cellulase EngZ for *Clostridium cellulovorans*, NucA from *Staphylococcus aureus*, and PelA_hyd_ from *P. aeruginosa*.

We validated the expression and secretion of the biofilm degrading enzymes from *L. plantarum* and LGG using SDS-PAGE of induced culture supernatants and enzymes activity assays. LAB cultures that contained NucA, EngH, or PelA_hyd_ expression vectors had protein bands and/or enzymatic activity in the filtered supernatants, which indicates successful secretion of the intended enzymes. Specifically, the supernatants of NucA- and PelA_hyd_-expressing LAB contained protein bands of the appropriate size (Fig. 3A) but we saw no visible band for EngH. The larger molecular weight of EngH compared to NucA and PelA_hyd_ puts it in a region where numerous other protein bands in the gel make it difficult to resolve individual proteins, so we also checked for enzymatic activity. We confirmed that the supernatants of LAB secreting EngH had CMCase activity (Fig. 3B) whereas the supernatants of LAB secreting NucA had DNase activity (Fig. 3C).

**Figure 3:**
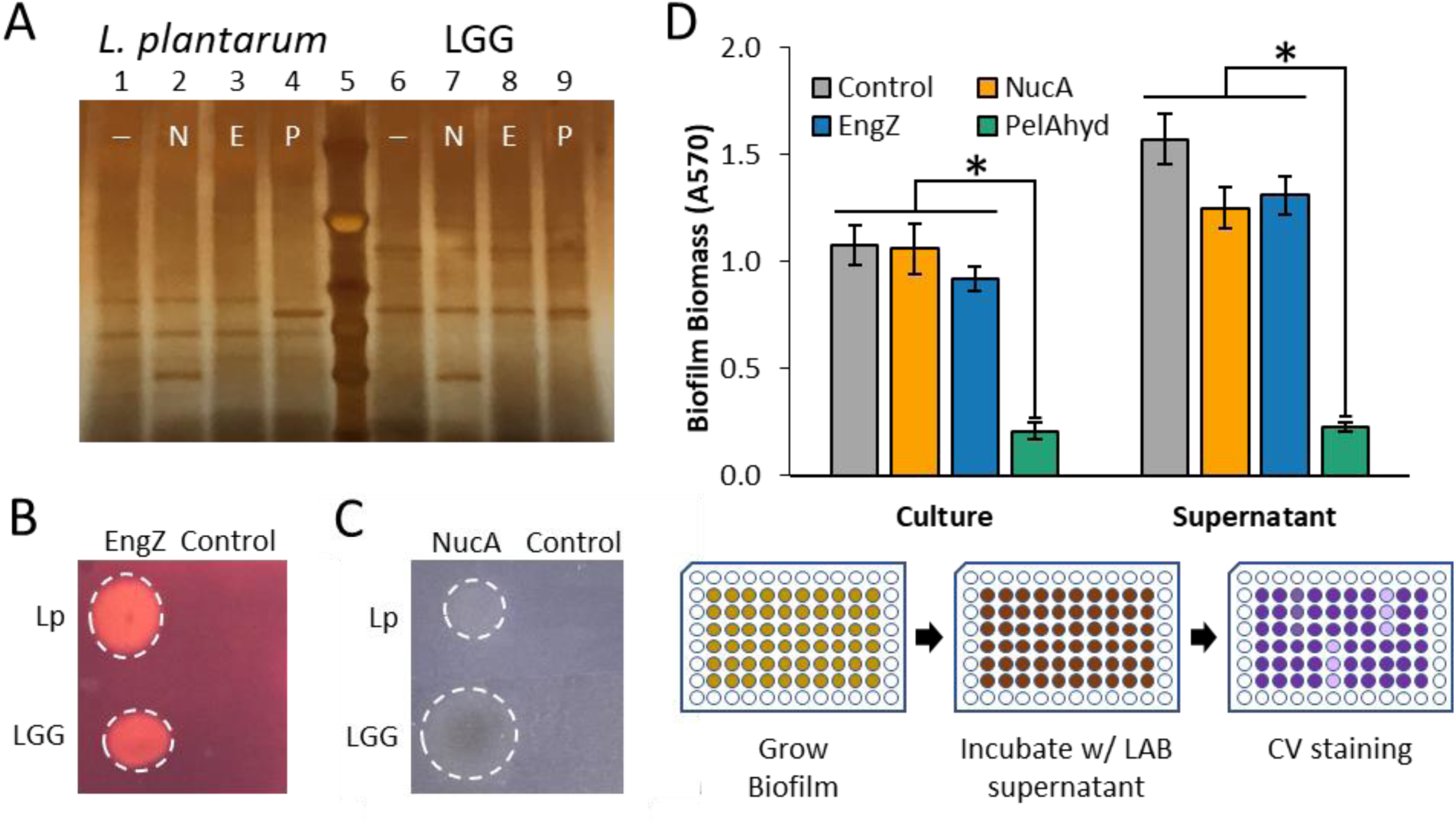
LAB-secreted enzymes degrade PA14 biofilm. (A) Silver stained SDS-PAGE gel of supernatants from induced LAB cultures; Lanes 1-4 *L. plantarum* containing control (empty vector pTCC210, ─), NucA (N), EngZ (E), and PelA_hyd_ (P) plasmids; Lane 5 Ladder; Lanes 6-9 LGG containing control (─), NucA (N), EngZ (E), and PelA_hyd_ (P) plasmids. (B) CMCase plate assay of EngZ-expressing *L. plantarum* and LGG. (C) DNase plate assay of NucA expressing *L. plantarum* and LGG. (D) Degradation of PA14 biofilms with the cultures and supernatants of *L. plantarum* containing control, NucA, EngZ, and PelA_hyd_ expression plasmids. Workflow of (D) is given below histogram. * denotes significant difference as determined by One-Way ANOVA (α 0.05) and Tukey HSD comparing samples of same type (e.g. cultures or supernatants); p < 0.01. Error bars represent ± 1 standard error. *n* = 12 for all conditions from 4 biological replicates.

We tested the ability of LAB cultures expressing and secreting NucA, EngH, or PelA_hyd_, as well as their cell-free (filtered) supernatants, to degrade mature PA14 biofilms. We chose to induce the cultures in BHI to decouple the growth and biofilm formation inhibition that we previously characterized using LAB cultures when grown in MRS, from the biofilm degradation capacity of the secreted enzymes. Cultures and supernatants of *L. plantarum* expressing PelA_hyd_ were highly effective at biofilm degradation, resulting in 80 % and 85 % reduction in biofilm biomass, respectively (Fig. 3D). However, EngZ and NucA expressing cultures and supernatants were ineffective at degrading PA14 biofilms. When we applied induced LGG cultures expressing the same proteins to PA14 biofilms, there was a considerable increase in biofilm biomass (Fig. S2). We were unable to conclude whether the engineered LGG cultures degraded PA14 biofilms due to this large increase in biofilm biomass. We did not observe an increase in biofilm biomass when we applied the filtered LGG supernatant to the PA14 biofilms, indicating that the increase in biomass was likely due to adhesion, growth, or biofilm formation by LGG itself. No biofilm was present when LGG was cultured in wells that did not contain PA14 biofilms, suggesting that the PA14 may aid in LGG surface adhesion.

### 3.4. Culture pH determines effectiveness of engineered *L. plantarum* anti-biofilm activity

Having established the significance of pH for PA14 growth and viability when treated with LAB cultures and supernatants, and knowing the optimal pH for NucA and EngZ are 9─10 and ∼7 ^39,40^, respectively, we postulated that we could enhance biofilm degradation by NucA and EngZ by modulating the supernatant pH. However, increasing culture pH did not significantly enhance biofilm degradation by NucA or EngZ relative to the control (Fig. 4A). We found no biofilm degradation by any of the enzymes when supernatants were buffered to pH 4.0 or pH 9.0. Interestingly, but unsurprisingly, formation of biofilm biomass was dramatically enhanced at pH 7.0 for all supernatants, although PelA_hyd_ was still effective at lowering biofilm biomass by 40 % relative to the control. NucA also moderately decreased PA14 biofilm biomass at pH 7.0, but the difference was not found to be significant (p > 0.05).

While performing the biofilm degradation assay, it became apparent that the large increase in CV staining at pH 7.0 and 9.0 was due to formation of additional PA14 biofilm. We illustrated this additional biofilm formation in test tubes (Fig. 4B). Mature biofilm treated with empty vector supernatant had a single biofilm at the air-solid-liquid interface at the original height of the culture volume during biofilm formation. When the control supernatant solution was buffered to pH 7.0, and added to the mature biofilm, a second biofilm formed at the height of the control supernatant, accounting for the higher biofilm biomass detected in the microplate assay. This second biofilm was not present when the mature biofilm was incubated with PBS, which reveals that PA14 can utilize residual nutrients in the *L. plantarum* supernatant to form additional biofilm. The unmodified and pH 7.0 buffered PelA_hyd_ supernatants degraded the mature biofilm, however some minimal new biofilm was formed when the pH was adjusted to pH 7.0.

**Figure 4:**
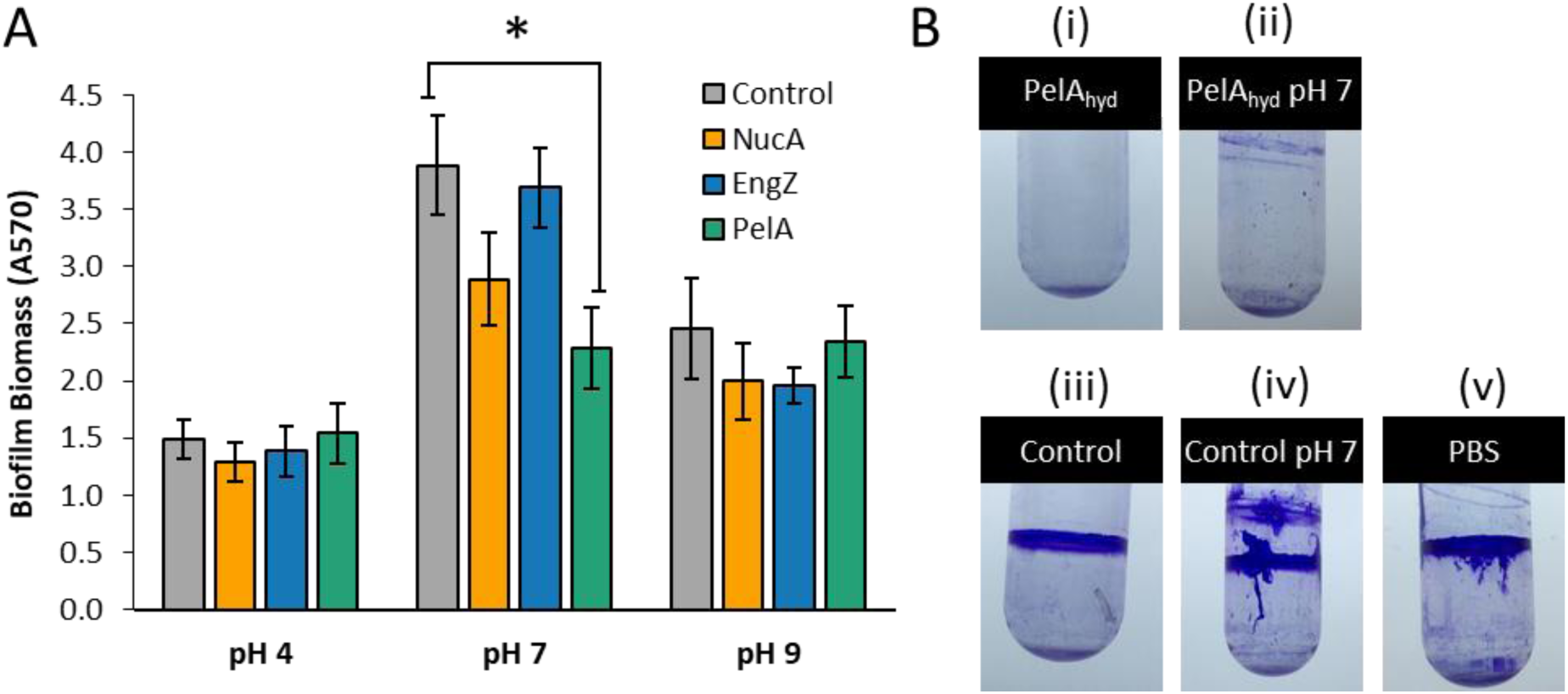
Effect of enzyme secreting *L. plantarum* culture supernatants on PA14 biofilm. (A) Degradation of PA14 biofilms with supernatants of *L. plantarum* containing control, NucA, EngZ, and PelA_hyd_ plasmids buffered to pH 4.0, pH 7.0, and pH 9.0. Workflow is the same given in Figure 3. * denotes significant difference as determined by One-way ANOVA (α 0.05) and Tukey HSD comparing samples of same pH; p < 0.05. Error bars represent ± 1 standard error. (B) Qualitative analysis of PA14 biofilms formed in culture tubes and treated with (i) PelA_hyd_ expressing *L. plantarum* supernatant with pH (i) unadjusted and (ii) adjusted to 7.0; supernatant from *L. plantarum* containing empty vector with pH (iii) unadjusted and (iv) adjusted to 7.0; (v) PBS.

## 4. Discussion

LAB effect their antimicrobial activity though a variety of mechanisms, including the production of antimicrobial proteins/peptides, inhibitory metabolites, and organic acids. While *L. plantarum* WCFS1 produces three bacteriocins (plantaricins A, EF, and JK), all of which act against a relatively narrow range of physiologically similar Gram-positives^41,42^, and *L. rhamnosus* GG produces an array of antimicrobial peptides, with varying degrees of activity against both Gram-positive and Gram-negative bacteria^43^, we found the distinguishing inhibitory factor of LAB supernatants and cultures against PA14 growth to be low pH. The inhibitory activity of both *L. plantarum* and LGG inversely correlated to decreasing pH and was abolished if the LAB supernatant was buffered to a more neutral pH. As heterofermenters, we expect both lactobacilli to produce lactic acid and acetic acid as fermentation products^44,45^. Though both of these acids inhibit the growth of Gram-negative pathogens like *P. aeruginosa*^46–48^, PA14 growth inhibition was indistinguishable from growth medium buffered to an equivalent pH range, indicating that the identity of the acid was not very important. The low pH of the LAB supernatants was also important for decreasing biofilm formation and biofilm viability, and was a major factor in the success of the degradation of biofilm by engineered LAB. Buffering the supernatants of LAB cultures directly or by dilution in PBS to more neutral pH resulted in maintenance of biofilm viability and the capacity to form new or more biofilms. Indeed, other investigations into multispecies bacterial communities have revealed that pH is a major determinant of success and failure within the community, and can generate a competitive advantage that results in elimination of acid-intolerant species^49^.

As a topical wound therapy, maintenance of the low pH would be beneficial for pathogen load reduction and clearance. Dilute organic acid solutions (e.g. acetic acid) have been used as a first-aid measure to stave off infection, yet we found that LAB cultures were more inhibitory than their acidic supernatants alone. Valdez et al. also found that *L. plantarum* cultures are more inhibitory to *P. aeruginosa* than the supernatants alone^50^. *L. plantarum* is known to remain viable at pH values below those found following 24 h culture in MRS (pH < 4), which would explain why cultures are better at inhibiting PA14 compared to supernatants; *L. plantarum* can continue to acidify, and increase inhibitory activity, by utilizing residual nutrients in the media or from the LB agar plates onto which they were applied. This illustrates a major advantage to the use of LAB cultures over acidified solutions or supernatants. LAB cultures can continue to acidify their environment when given additional nutrients, thus maintaining an antimicrobial environment. Similarly, LAB cultures could conceivably continue to consume fermentable substrates in the wound bed, competing with pathogens for nutrients. In fact, the skin normally maintains an acidic pH (4.0 ─ 5.5), and the normal wound healing processes—including decreased metalloprotease activity and epithelial migration—are correlated with decreasing pH. Conversely, elevated wound pH is often associated with chronic wounds. Thus, LAB-mediated acidification should create an inhospitable environment for acid-intolerant pathogens and is not expected to have a negative impact on the normal wound healing process.

Interestingly, we found that even though the two lactobacilli investigated in this study – *L. plantarum* and LGG – inhibited PA14 viability by lowering the culture pH, their outcomes were divergent when applied to biofilms. The considerable increase in biofilm biomass seen following the addition of LGG culture to PA14 biofilms was dependent upon the presence of an extant matrix, suggesting LGG adheres to PA14 cells or EPS. While LGG is known to produce adherent biofilms, it is not known to do this when grown on MRS^51^, which we found as well. While we assume that the PA14 present in this co-culture biomass is no longer metabolically active because LGG supernatants rendered PA14 biofilms non-viable, we elected to disregard LGG as a potential bacteriotherapy as our stated intent was to decrease biofilm biomass. However, it remains entirely possible that the increased adhesion of LGG could enhance its antimicrobial effect by maintaining close proximity to the pathogen —and this interaction may be worth of future investigations.

After engineering the lactobacilli to secrete a series of biofilm degrading enzymes, we found that only PelA_hyd_ secreted by *L. plantarum* was effective at degrading PA14 biofilms. Surprisingly, DNAse and EngZ secreted by *L. plantarum* were unable to appreciably degrade PA14 biofilms even though previous investigations have shown the efficacy of enzymes of these classes to be effective anti-biofilm agents against this strain. We verified the secretion and activity of NucA and EngZ in the LAB supernatant and optimized the supernatant pH for the activity of these enzymes, and still found no significant benefit. The activity of these enzymes at elevated pH may be masked by the additional growth of PA14 biofilm at the elevated pH at which these enzymes are most active. However, EngZ exhibits approximately 60% activity even at pH 4.0, and yet we still saw no impact on PA14 biofilm degradation. Previous work has shown that cellulases extracts from *Trichoderma viride* or *Aspergillus niger* can degrade PA14 biofilms^37,52^. However, the biofilm degrading capacity was not attributed to any single enzyme or endoglucanase activity and the exact composition of the extract is unknown. Further, activity on Pel is likely due to substrate promiscuity, which is often enzyme dependent. Thus, EngZ may not have the same range of relaxed substrate specificity as *T. viride* or *A. niger* cellulases. The differences in our observations compared to that in literature could also be due to differences is assay conditions. Specifically, PA14 biofilms degradation by DNase was shown in flow cells, where DNA is known to play an integral role in the structure of biofilm stalks at the solid-liquid interface when under flow^52,53^. DNA may not play the same role in static batch cultures where the biofilm forms at the air-solid-liquid interface. Additionally, the DNA present in flow cell biofilms only plays an important adhesive role in early stage attachment^36^, and may not play a critical role in maintenance of mature biofilms. DNA also contributes to a plethora of interesting phenotypes in the biofilm, including chelating cations, inducing antibiotic resistance, promoting inflammation, and aiding extracellular electron transport, all of which are important metrics by which to test this therapy in the future^54–57^.

Through the development of this bacteriotherapy for the disruption of PA14 biofilm, we learned potentially important design rules for engineered bacteriotherapies. Selection of an appropriate organism as the chassis to engineer for the bacteriotherapy is important. Specifically, the bacteriotherapeutic organism’s ability to inhibit the pathogen growth, biofilm formation and impact on mature biofilms, using the assays described in this work, can determine whether the organism will act as an effective bacteriotherapy. We found that prevention of additional biofilm formation and pathogen growth is a key to the degradation process. Additionally, the selection of the appropriate enzyme for biofilm degradation is equally important. We found that PA14-derived PelA_hyd_ was most effective at degrading its own biofilm, and that other enzymatic activities thought to be effective at degrading PA14 biofilms were ineffective in our assay. Frequently, genes have been identified within the genome of biofilm-forming organisms that function to degrade the biofilm and release the embedded cells. However, their expression is often suppressed during the biofilm growth phase to ensure biofilm integrity. Therefore, we propose that the enzymes for biofilm degradation should be sourced from the pathogen itself, as these native enzymes were “designed” to degrade the EPS polymers. Such observations are consistent with previous studies.

As current antimicrobial treatments decrease in efficacy, the development of novel treatments is essential for effectively treating recalcitrant infections. Traditional small molecule screening for antimicrobials has all but ceased due to the high cost and uncertainty of success. Engineered bacteriotherapies provide an alternative strategy for developing antimicrobials, with specific component parts (organism, enzyme, intended pathogen) that can be intentionally modified to address the challenges of particular infections. Though we present promising data for the ability to target *P. aeruginosa* PA14 biofilms using an engineered bacteriotherapy, further validation of this system is required *in vivo*. Expanding the pathogen targets, host infection sites, and adding additional functionalities, such as the production of specific antimicrobials, will better validate this system as an effective treatment alternative to existing therapies.

## 5. Acknowledgements

We would like to thank Dr. David R. Snydman (Tufts Medical Center, Boston, MA), Dr. Abraham L. Sonenshein (Tufts University School of Medicine, Boston, MA), Dr. Michiel Kleerebezem (Wageningen University & Research, Wageningen, Netherlands), Dr. Roberto Kolter (Harvard Medical School, Boston, MA), Dr. Ann Hochschild (Harvard Medical School, Boston, MA), Dr. Huimin Zhao (University of Illinois, Urbana, IL), Dr. Geir Mathiesen (Norwegian University of Life Sciences, Ås, Norway), and Dr. Jan Peter van Pijkeren (University of Wisconsin, Madison, WI) for graciously providing strains and/or plasmids required to complete this work.

