## Supplemental Information for "Engineered lactobacilli display anti-biofilm and growth suppressing activities against *Pseudomonas aeruginosa*"

4 Colby St, STC 276

Medford, MA 02155

617-627-2582

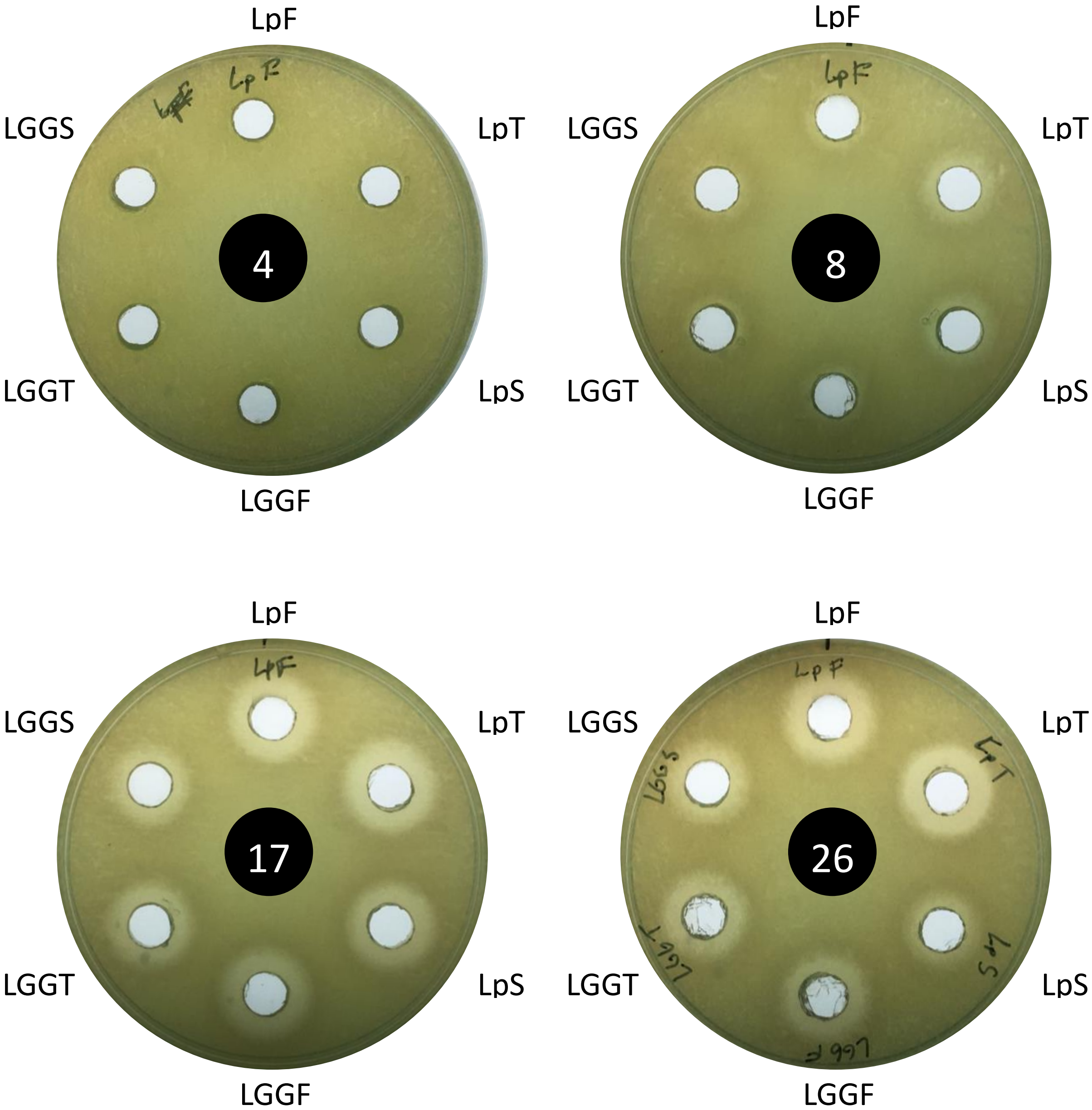


Fig. S1: Impact of LAB culture aeration and duration on PA14 growth inhibition. *L. plantarum* (Lp) and *L. rhamnosus* (LGG) were grown shaking in a baffled flask (F), shaking in a test tube (T), or statically in a test tube (S) for 4, 8, 17, or 26 hours.

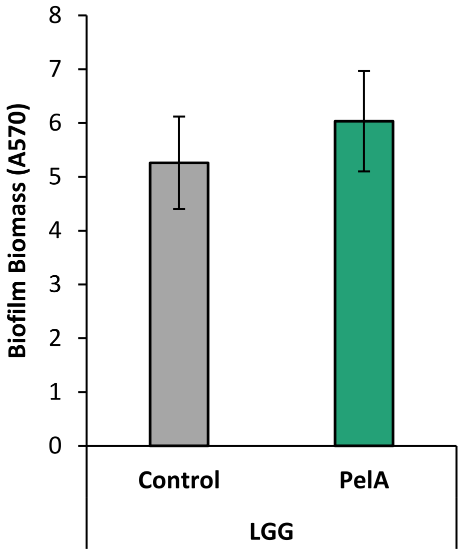


Fig. S2: PA14 biofilms treated with LGG cultures containing the Control or $PelA_{hyd}$ plasmid.

**Table S1:** Bacterial strains used in this study.

| Strain | Genotype/Use/Origin | Source/Origin |
|---|---|---|
| *Lactobacillus rhamnosus* GG | Human stool isolate [1] | Dr. David R. Snydman, TMC |
| *Lactobacillus plantarum* WCFS1 | Single colony isolate from NCIMB8826, an isolate of human saliva[2] | Dr. Michiel Kleerebezem, WUR |
| *Pseudomonas aeruginosa* PA14 | Clinical isolate from burn wound[3] | Dr. Roberto Kolter, HMS |
| *E. coli* TG1 | cloning | Dr. Ann Hochschild, HMS |
| *E. coli* TG1 *endA*$^{-}$ | *endA::cat*; cloning | This work |
| *E. coli* DH5a Z1 | Cloning; R. Lutz and H. Bujard 1997[4] | Dr. Huimin Zhao, UIUC |
| *Staphylococcus aureus* UAMS-1 | Source of *nucA*. | Dr. Abraham L. Sonenshein, TUSM |
| *Clostridium cellulovorans* DSM 3052 | Genomic DNA. Source of EngZ | DSMZ |

**Table S2:** Plasmids used in this study.

| Plasmid | Notes | Source |
|---|---|---|
| pLp_3050Ag85B-E6cwa2 | Expression vector for *L. plantarum* derived from pSIP401[5]. low copy narrow host 256rep ori + pUC ori. Expression induced by addition of SppIP. | Dr. Geir Mathiesen, NMBU |
| pSIP411 | Broad host, High copy SH71 ori only. LAB expression vector.[6] | Dr. Jan Peter van Pijkeren, UW |
| pTCC200 | pSIP401 with a new multiple cloning site, N-terminal 6x-histidine tag, and Lp_3050 secretion signal | This work |
| pTCC204 | pSIP401 with *engZ* | This work |
| pTCC210 | pSIP411 with a new multiple cloning site, N-terminal 6x-histidine tag, and Lp_3050 secretion signal | This work |
| pTCC211 | pTCC210 with *nucA* | This work |
| pTCC214 | pTCC210 with *engZ* | This work |
| pTCC216 | pTCC210 with *pelA*$_{hyd}$ | This work |

**Table S3:** Primers used in this study.

| # | Name | Sequence |
|---|---|---|
| 1 | Pro1 | gactcagatctaccggtttaatttgaaaattg |
| 2 | Pro2 | taaaatctccttgtaatagtattttatagaatac |
| 3 | Pro3 | ctataaaatactattacaaggagattttacatATGAAAAAATTTAACTTTAAAACC |
| 4 | Pro4 | CCGTGGAACTAAACCTGATGAATGATGATGATGATGATGCGTACGCTTGGAGGCCTGGGC |
| 5 | Pro5 | gcactcacgtgccatggcatgcgtcgacTGAACCCCGTGGAACTAAACCTGATGAATG |
| 6 | nucAF | gactcgtcgactcaactaaaaaattacataaagaacc |
| 7 | nucAR | gcactcacgtgttattgacctgaatcagcgttg |
| 8 | engZF | gactcgtcgacacagaaaattacaactacgggg |
| 9 | engZR | gcactcacgtgttagaaactagttatttgacctaaaatgtattttttc |
| 10 | pelAhydF | gtttagttccacggggttcaGGCGGGCCGTCCAGCGTG |
| 11 | pelAhydR | gtgccatggcatgcgtcgacTCACGGTTGCACCTCGACGTCG |
| 12 | 411BuilderF | GTCGACGCATGCCATGGC |
| 13 | 411BuilderR | TGAACCCCGTGGAACTAAACC |
| 14 | pSIPseqF | CAGCTCCAGATCTACCGG |
| 15 | pSIPseqR | gcaatatcagtaattgctttatcaactgctg |
| 16 | engZmidseqF | GAGCTTTGCTTAAAGCACG |
| 17 | endAKOF | CTTTCGCTACGTTGCTGGCTCGTTTTAACACGGAGTAAGTGGCTGCTTCGAAGTTCC |
| 18 | endAKOR | CTGGCCTTCACCGCCATTCCACTGGCTGTACATAAAGTTGCCTCCTTAGTTCCTATT |
| 19 | endAverF | cctgatctggctgattgcatacc |
| 20 | endAverR | ctcccagtcggtaaccggatac |
